# Computational Modeling of Effects of *PKP2* Gene Therapy on Ventricular Conduction Properties in Arrhythmogenic Cardiomyopathy

**DOI:** 10.1101/2024.12.12.628155

**Authors:** Alessio Ostini, André G. Kléber, Yoram Rudy, Jeffrey E. Saffitz, Jan P. Kucera

**Author notes:** **Corresponding author:** Jeffrey E. Saffitz, MD, PhD, Department of Pathology, Beth Israel Deaconess Medical Center, 99 Brookline Avenue, Boston, MA 02215 USA. These authors contributed equally (Jeffrey E. Saffitz, Jan P. Kucera).

## Abstract

**Background:** Patients with arrhythmogenic cardiomyopathy (ACM) due to pathogenic variants in *PKP2*, the gene for the desmosomal protein plakophilin-2, are being enrolled in gene therapy trials designed to replace the defective allele via adeno-associated viral (AAV) transduction of cardiac myocytes. Evidence from experimental systems and patients indicates that ventricular myocytes in *PKP2* ACM have greatly reduced electrical coupling at gap junctions and reduced Na^+^ current density. In previous AAV gene therapy trials, <50% of ventricular myocytes have generally been transduced.

**Methods:** We used established computational models of ventricular cell electrophysiology to define the effects of varying levels of successful gene therapy on conduction in patients with *PKP2* ACM. Conduction velocity and development of conduction block were analyzed in tissue constructs composed of cells with levels of electrical coupling and Na^+^ current density observed in experimental studies.

**Results:** We observed a non-linear relationship between conduction velocity and the proportion of transduced cells. Conduction velocity increased only modestly when up to 40% of myocytes were transduced. Conduction block did not occur in tissue constructs with moderate levels of uncoupling (0.10 or 0.15 of normal) as this degree of coupling was sufficient to allow electrotonic current to pass through diseased cells. Thus, low levels of transduction, likely to occur in phase 1 clinical trials, do not appear to pose a major safety concern. However, our models did not incorporate potential effects of fibrosis and immune signaling, both of which will presumably be present in *PKP2* ACM patients undergoing gene therapy.

**Conclusions:** The extent of successful ventricular myocyte transduction anticipated to be achieved in *PKP2* AAV gene therapy trials will likely not restore conduction velocity to levels sufficient to decrease risk of reentrant arrhythmias.

**What is Known:**

- Patients with arrhythmogenic cardiomyopathy due to pathogenic variants in *PKP2* (the gene for the desmosomal protein plakophilin-2) are now being enrolled in gene therapy trials.
- Experimental and clinical observations indicate that patients with arrhythmogenic cardiomyopathy have slow ventricular conduction with a propensity to conduction block due to source-sink mismatch.
- <50% of ventricular myocytes are usually transduced after adeno-associated viral gene therapy.

**What the Study Adds:**

- At anticipated levels of successful transduction of ventricular myocytes, little change in conduction velocity will be achieved in patients with arrhythmogenic cardiomyopathy due to variants in *PKP2*.
- Higher levels of transduction could produce conditions that increase risk of conduction block, especially in the presence of areas of non-conducting fibrofatty scar tissue.

## Introduction

Three companies are currently enrolling patients with arrhythmogenic cardiomyopathy (ACM) caused by pathogenic variants in *PKP2* (the gene for the desmosomal protein plakophlin-2) to undergo gene therapy via adeno-associated viral (AAV) transduction of cardiac myocytes.^1-3^ Patients with active ACM due to *PKP2* variants typically manifest features of classical arrhythmogenic right ventricular cardiomyopathy including ventricular arrhythmias with left bundle branch morphology, progressive usually right ventricular degeneration with accumulation of fibrofatty scar tissue, and increased risk of adverse events caused by exercise.^4-6^ The goal of these gene therapy trials is to replace the defective *PKP2* allele with a normal gene and thereby mitigate clinical features of disease and reduce risk of sudden death.

Several studies have shown that ventricular myocytes that express pathogenic ACM variants have reduced immunoreactive signal for connexin43 (Cx43), the major ventricular gap junction protein, at intercalated disks, along with marked reduction in sodium current (I_Na_) density and modest resting membrane depolarization due to reduced inwardly rectifying potassium current (I_K1_) density.^7-9^ Our recent in vitro study reported ∼65% reduction in conduction velocity in strands of ventricular myocytes that expressed a pathogenic *PKP2* variant associated with increased rate-dependent development of unidirectional conduction block due to source-sink mismatch.^10^ As determined by computer modeling, these conduction abnormalities were largely attributable to ∼85% reduction in electrical coupling between cells (associated with greatly reduced junctional Cx43 signal) and ∼50% reduction in peak I_Na._^10^ Reduced immunoreactive signals for Cx43 and Na_v_1.5 at ventricular intercalated disks have been reported in patients with *PKP2* ACM ^11-13^ and electrocardiographic imaging (ECGI) has demonstrated slow non-uniform conduction and fractionated electrograms in *PKP2* ACM patients.^14^Considered together, these studies indicate that conduction in ventricular myocardium in patients with active *PKP2* ACM will likely be slow due to reduced electrical coupling and reduced I_Na_ density in ventricular myocytes. To define the potential effects of gene therapy on ventricular conduction in patients with *PKP2* ACM, we performed computer simulations using two models of ventricular myocyte electrophysiology in tissue constructs composed of cells with decreased levels of electrical coupling and I_Na_ density as observed in experimental studies. We assessed the effects of varying degrees of transduction by AAV (0-100% success in replacing the defective gene) on conduction velocity and the propensity to develop conduction block. Our results show that little change in conduction velocity is achieved even after the electrical phenotype of 40-50% of cells is corrected.

## Methods

Conduction was simulated in rectangular strands of tissue composed of excitable elements (60 × 60μm cells) arranged on a square lattice (Figure 1A). The strands were 180 elements long (1.08 cm). Strands of two different widths were analyzed: wide strands (21 elements across; 1.26 mm) and narrow strands (5 elements across, 0.3 mm).

**Figure 1.**
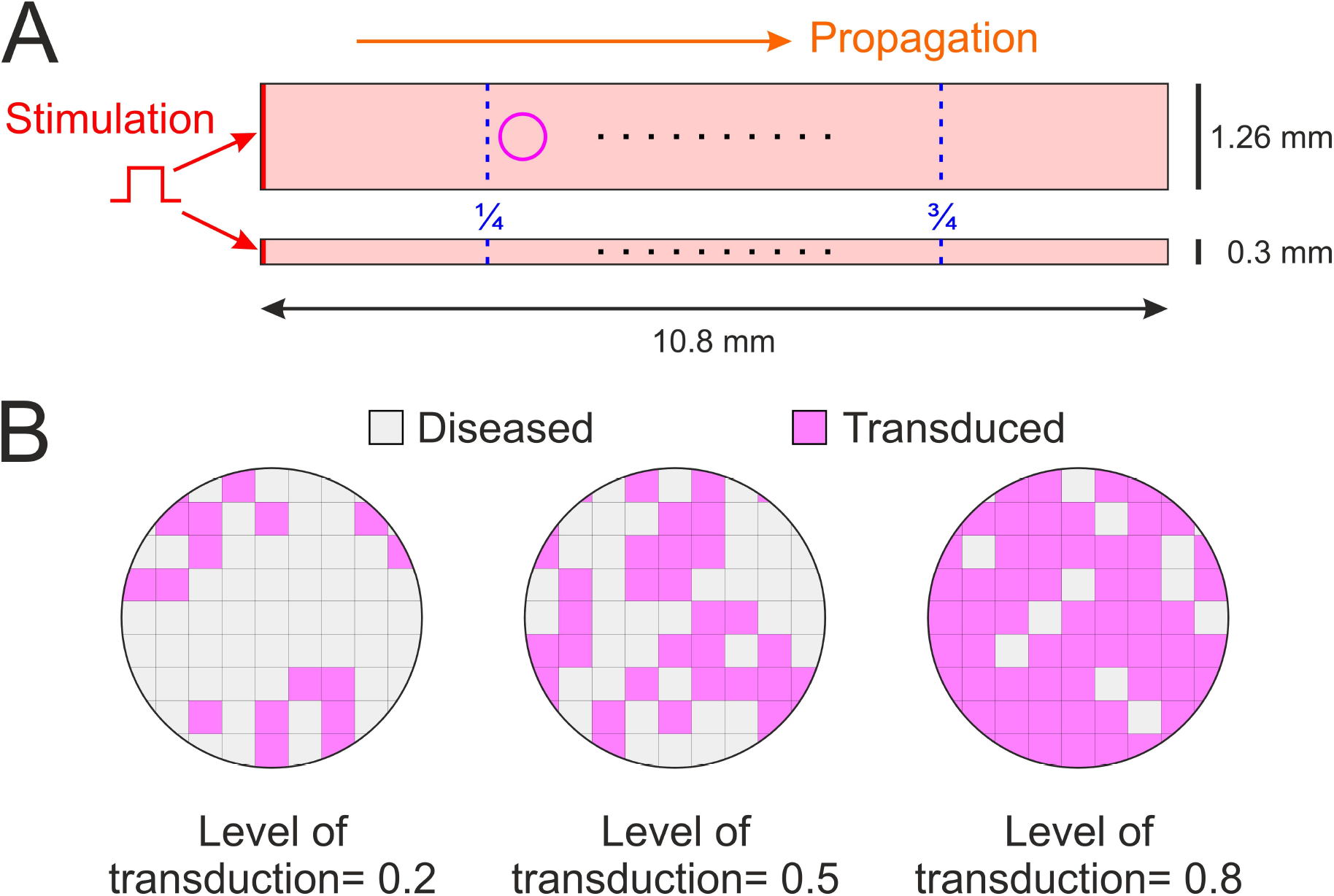
Schematic representation of the simulation setup. **A**: Cardiomyocyte strands were represented as rectangular domains (top: wide strand; bottom: narrow strand). A stimulus was applied to the left edge (red) to trigger action potential propagation (orange). The magenta circle indicates the region magnified in B. Black dots indicate sites where the action potential waveform was registered. **B**: Illustration of the simulated cellular arrangement corresponding to the circle in **A**. The square lattice represents the 2-dimensional tissue model in which each square corresponds to an individual cell. Grey cells represent diseased *PKP2* ACM cells, while magenta cells represent transduced cells (normalized to a wild-type phenotype) randomly positioned in the network at predefined levels of transduction (left: 0.2; middle: 0.5; right: 0.8).

To model membrane ionic currents, we used the classical Luo-Rudy phase I model ^15^ (LR1 model) and a modified LR1 model developed in previous studies to replicate experimental findings in cultures of cardiac myocytes from small rodents.^10,16-18^ These modifications to the LR1 model are described in detail in De Waard et al. (DeW model).^18^ Compared to the LR1 model, the main differences in the DeW model pertinent to propagation are i) lower maximum sodium current conductance (to account for slower action potential upstrokes in cultured myocytes), ii) increased slow inward calcium current with accelerated kinetics, and iii) reduced inward rectifier potassium current.^18^ The LR1 model and its DeW adaptation have been well-characterized in simulating cardiac conduction in the process studied here which depends primarily on the sodium current and intercellular coupling.^18,19^

To simulate ventricular myocytes in *PKP2*-ACM, the maximal sodium current conductance was reduced to 0.5, 0.4 and 0.3 times its normal value (i.e., the standard value in the LR1 or DeW model for wild-type (WT) healthy myocardium), in agreement with experimental measurements.^10^ To simulate decreased intercellular coupling in *PKP2*-ACM ventricular myocytes, the coupling conductance between an affected *PKP2*-ACM cell and any other cell (either a transduced or non-transduced *PKP2* ACM cell) was reduced to 0.15 or 0.1 times the normal value between two WT cells. These scaling values are in agreement with previous experimental observations,^10^ and conform to the notion that the coupling level between two connected cells is dictated by the cell expressing the smallest amount of gap junction protein at the cell-cell junction. In additional simulations, we examined the effects of reducing this coupling level further to 0.05 of normal. At a macroscopic level, the simulated tissues were isotropic.

Negative control simulations were performed in strands composed entirely of *PKP2* ACM cells. Varying degrees of successful gene transduction levels were simulated by randomly assigning cells as being either *PKP2*-ACM or transduced (WT), with the probability of being transduced ranging from 0.1 to 0.9 in steps of 0.1. This resulted in random mosaic patterns of the two cell types (Figure 1B). Positive control simulations were performed in strands composed of 100% transduced (WT) cells.

Propagation in the x-direction was initiated at the edge of the strands by injecting a supra-threshold current pulse into the first column of elements (Figure 1A). Conduction was simulated in 10 realizations of the random mosaic pattern for each combination of model (LR1 or DeW), strand width, sodium current conductance, level of coupling and level of transduction. Activation time was defined at –35 mV during depolarization. For successfully propagated impulses, conduction velocity was determined by linear regression of the earliest activation time at a given position (x) vs. x between ¼and ¾ of strand length to avoid sealed-end effects at the extremities of the strand. Propagation block was defined as failure of the wave front to reach ¾ of strand length.

Simulations were run with a constant time step of 0.005 ms. Membrane potential and intracellular calcium concentrations were integrated using the forward Euler method. Gating variables were integrated using the approach of Rush and Larsen.^20^ All simulations were conducted in MATLAB (version R2021a or higher, MathWorks).

## Results

### Partial transduction of cells causes heterogeneous conduction patterns

Conduction along a strand of 100% diseased *PKP2* ACM cells, simulated with the LR1 model, was slow (10.7 cm/s), but the propagation pattern was uniform and homogeneous, characterized by regular and similar action potential (AP) upstrokes, planar and regularly spaced isochrones, and a smooth activation profile (Figure 2A). Conduction through strands composed of 100% transduced (WT) cells was ∼5-times faster (52.4 cm/s) and was homogeneous as well (Figure 2B). By contrast, conduction velocity through tissue strands in which 40% of cells were transduced was only modesty increased above that seen in *PKP2* ACM strands (15.2 cm/s) and was characterized by irregular, dissimilar AP upstrokes and distorted isochrones (Figure 2C). This distortion of upstrokes and isochrones was caused by the heterogeneous levels of intercellular coupling and I_Na_, causing irregular conduction at the cellular level. This heterogeneity was also reflected by small deviations of the activation profile from a strictly linear pattern (see inset in Figure 2C).

**Figure 2.**
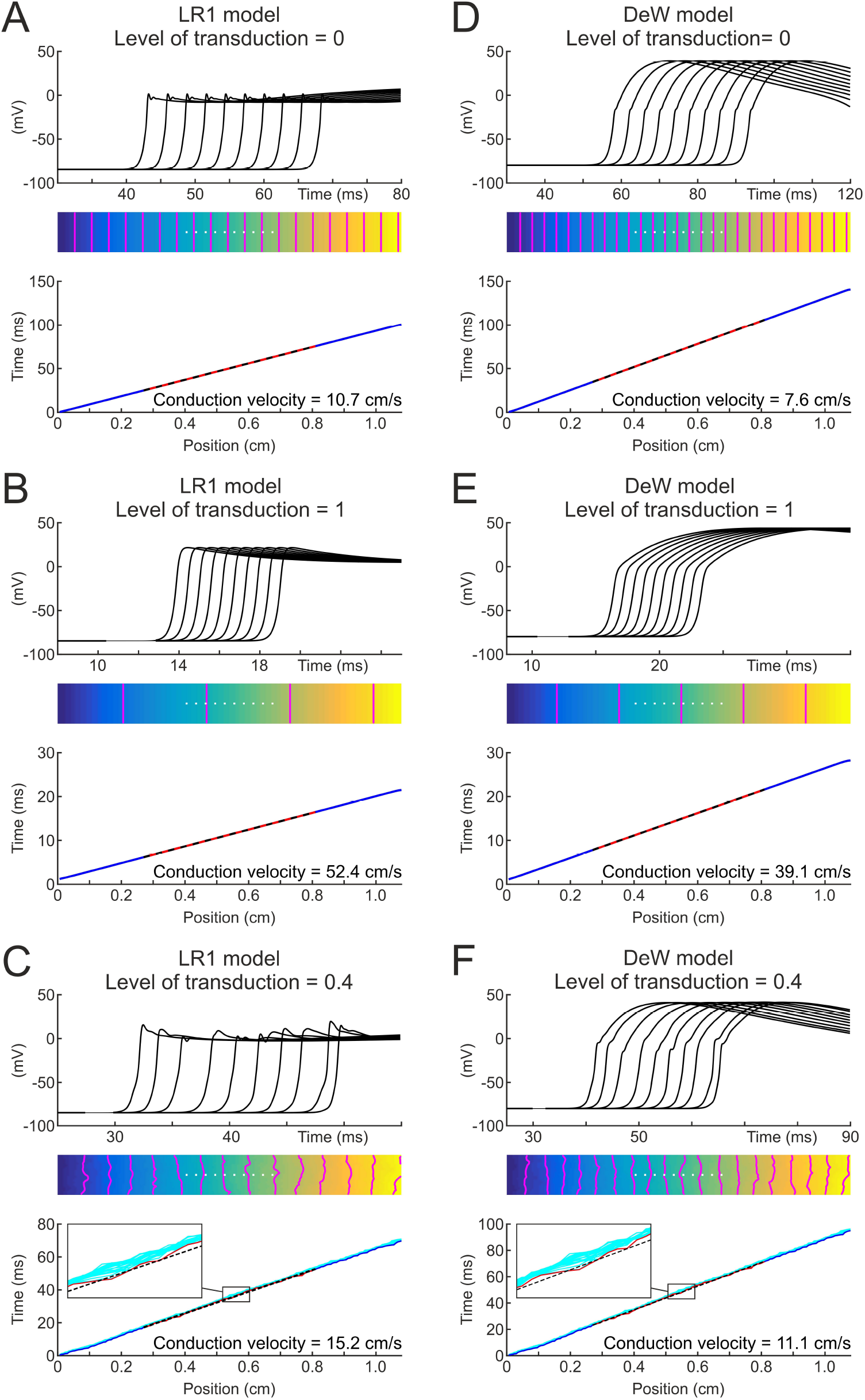
Simulated action potential propagation in wide strands under different conditions. **A**: Propagation in the LR1 model with 100% diseased cells (negative control; sodium current conductance reduced to 0.4 times its normal value; coupling to reduced to 0.1 times its normal value). Top: Action potential upstrokes at sites marked with white dots in the isochronal activation map below. Middle: Isochronal map (interval between isochrones (magenta): 5 ms). Bottom: Activation profile along the strand (blue: earliest activation time vs. position; red: earliest activation time used for linear fitting and conduction velocity determination; dotted black line: linear fit). **B**: Propagation in the LR1 model with 100% transduced cells (positive control); same layout as in A. **C**: Propagation in the LR1 model with 40% transduced cells (sodium current conductance reduced to 0.4 times normal value; coupling between cells reduced to 0.1 times its normal value); same layout as in A and B. Inset shows the dispersion of activation time (cyan lines), also reflected by the distorted isochrones. **D, E** and **F**: same as **A, B** and **C**, respectively, with the DeW model.

Qualitatively similar results were obtained using the DeW model. Conduction velocity was 7.6 cm/s in *PKP2* ACM cells (Figure 2D) and 39.1 cm/s in strands composed entirely of transduced (WT) cells (Figure 2E). Conduction velocity increased from 7.6 to 11.1 cm/s when 40% of cells were successfully transduced (Figure 2F).

### A low efficiency of transduction only marginally restores conduction velocity while higher efficiency may potentiate conduction block

We ran a series of simulations to examine the effects of different degrees of AAV transduction on conduction. Figure 3A shows conduction velocity in the wide LR1 strand model, in which different levels of I_Na_ conductance (0.5, 0.4 and 0.3 times that of control shown in blue, green and red, respectively) were assumed for the *PKP2* ACM phenotype with the same level of coupling (0.15 times the WT value) between a diseased cell and another cell. Ten simulations were run at each level of transduction (0.1 – 0.9) with different realizations of the random distribution of cells. At all three levels of I_Na_, the relationship between conduction velocity and transduction rate was non-linear, whereby an increase in transduction efficiency from 0 to 0.5 resulted in a relatively small restoration of conduction velocity compared to a further increase in efficiency from 0.5 to 1. The results were similar in the narrow LR1 strand model (Figure 3B). Qualitatively similar results were also seen in the DeW model (wide strand in Figure 3C; narrow strand in Figure 3D). Importantly, conduction block did not occur in this series of simulations.

**Figure 3.**
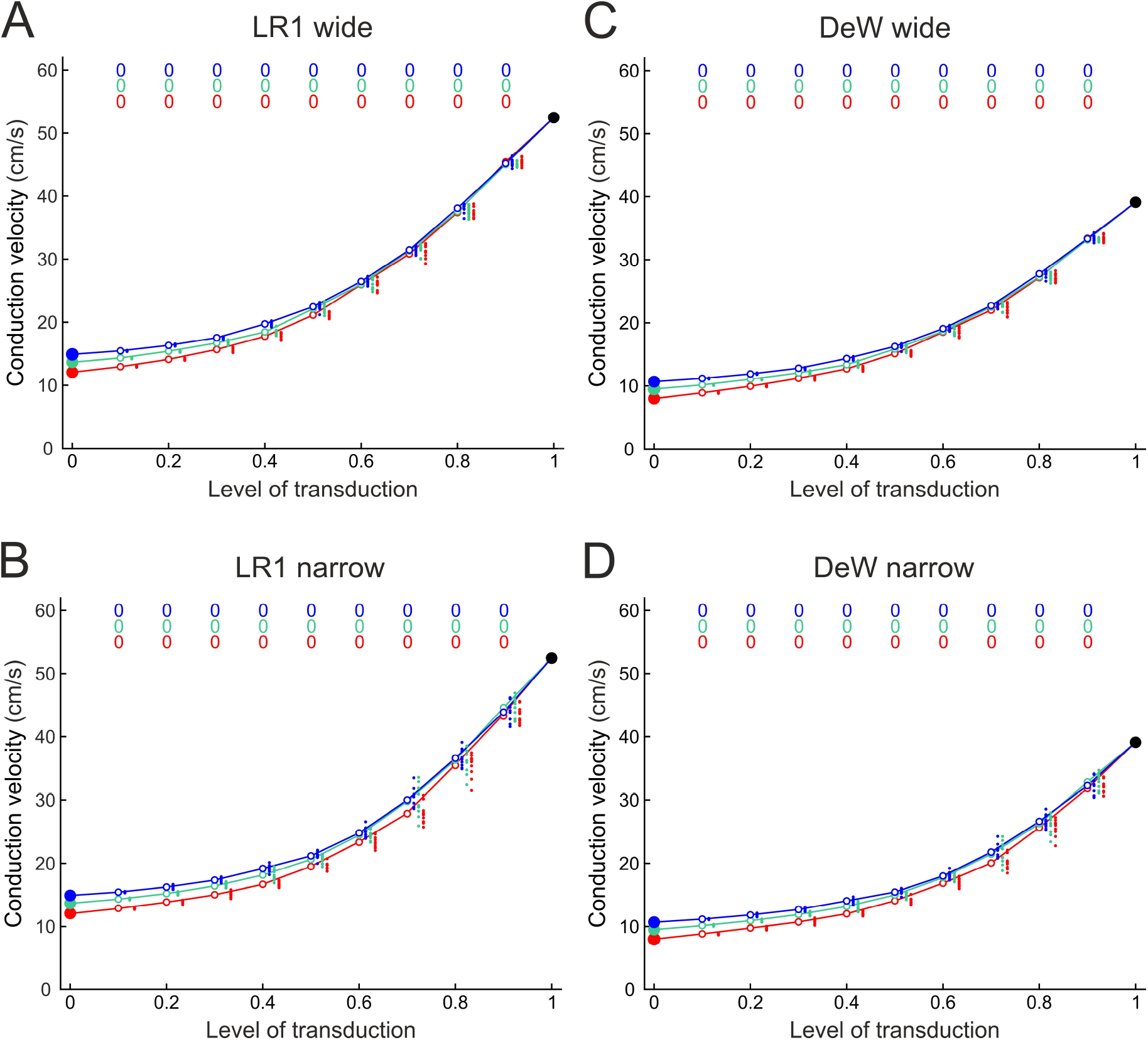
Effects of increasing transduction efficiency on conduction velocity when the coupling of a diseased cell with another cell was set to 0.15 times the normal value. Each panel shows conduction velocity vs. the proportion of transduced cells in the wide LR1 model strand (**A**), the narrow LR1 model strand (**B**), the wide DeW model strand (**C**) and the narrow DeW model strand (**D**). The sodium current conductance in diseased cells was reduced to 0.5 (blue), 0.4 (green) or 0.3 (red) of its normal value. Solid colored circles at zero transduction level indicate conduction velocity in untreated tissue. Solid black circles at full transduction level (1.0) indicate conduction velocity in normal (i.e., fully transduced) tissue. Open circles and connecting lines indicate the mean of conduction velocity in n=10 separate simulations with different random distributions of transduced cells. Individual data are shown as small colored dots. Colored numbers above the plots indicate the proportion of simulations in which conduction block occurred.

The same simulations were run at a lower coupling level (0.1 times the control value) (Figure 4). The results were again similar, but conduction block occurred in one simulation with the narrow LR1 model strand (Figure 4B). Finally, we ran a series at an even lower coupling level (0.05 of normal) (Figure 5). In general, we saw a similar non-linear dependence of conduction velocity on transduction efficiency, but in the narrow LR1 model strand conduction exhibited a substantial non-monotonic propensity to block at transduction rates from 0.4 to 0.9.

**Figure 4.**
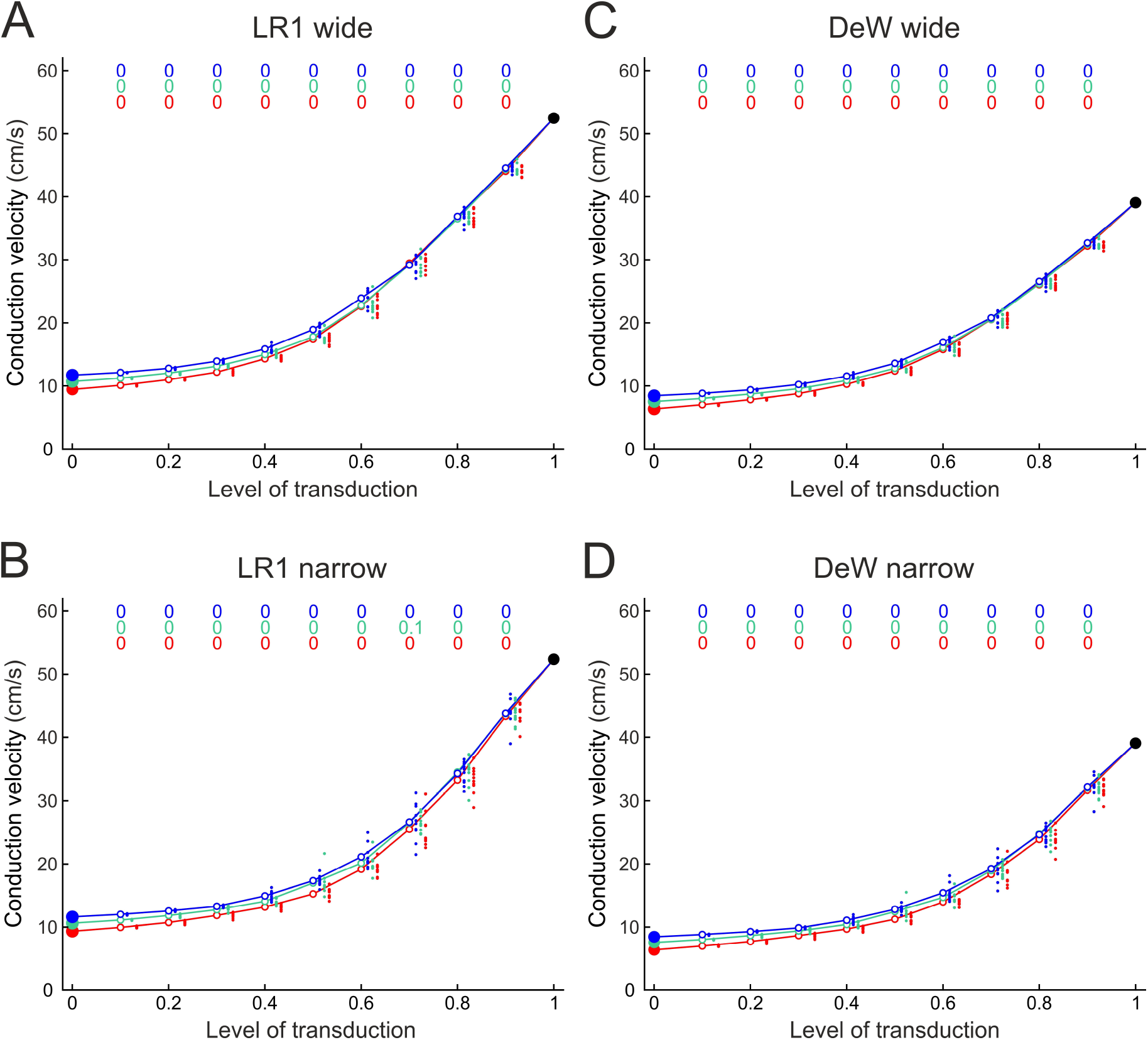
Effects of increasing treatment efficacy on conduction velocity when the coupling of a diseased cell with another cell was set to 0.1 times the normal value. Same layout as in **Figure 3**.

**Figure 5.**
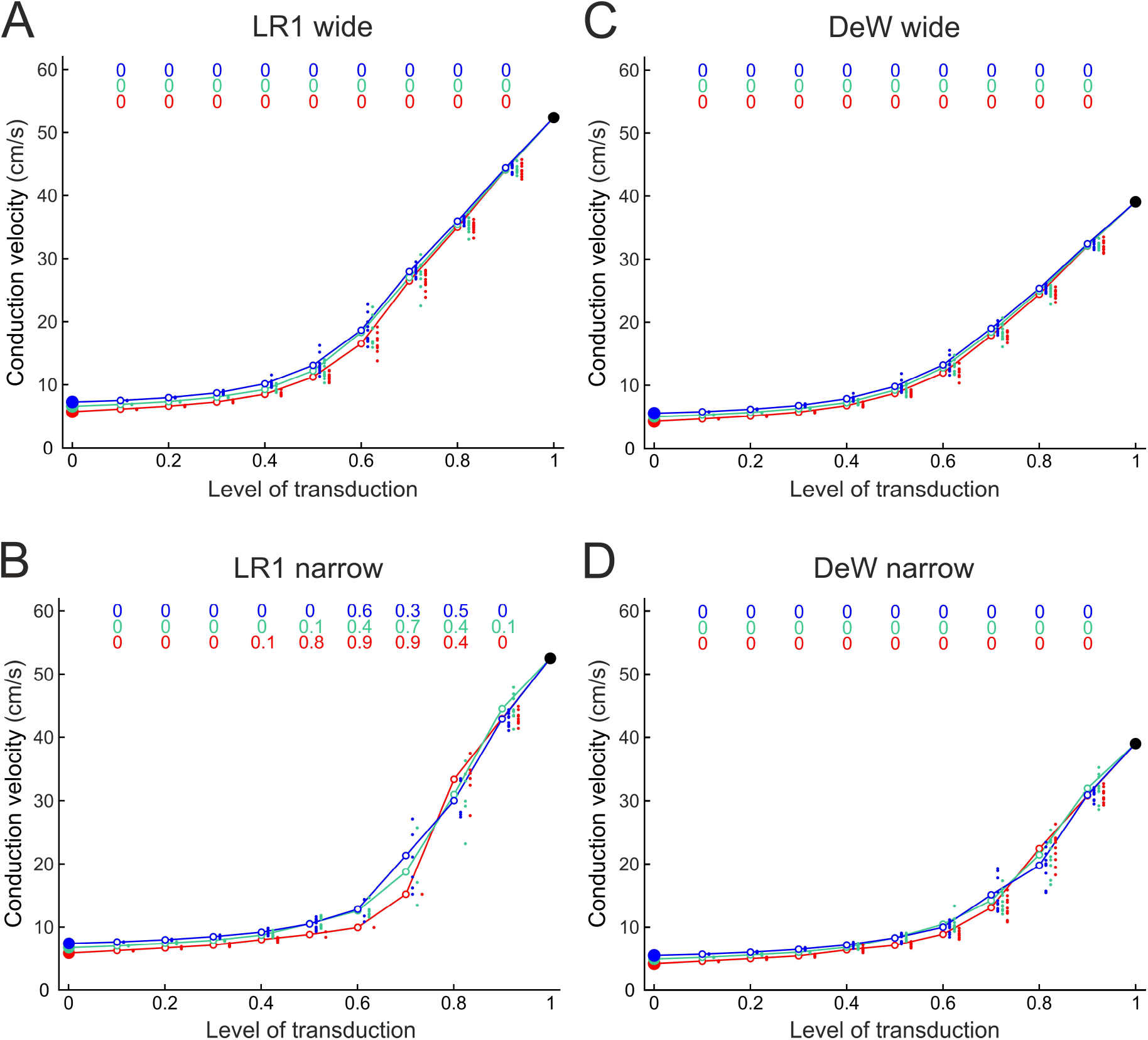
Effects of increasing treatment efficacy on conduction velocity when the coupling of a diseased cell with another cell was set to 0.05 times the normal value. Same layout as in **Figures 3** and **4**. Colored numbers above the plots indicate the proportion of simulations in which conduction block occurred. With this level of coupling, conduction block occurred in certain simulations, reflected by non-zero values above the plot in panel **B** (LR1 narrow strand model).

As summarized in Table 1, our results indicate that up to 40% successful viral transduction of ventricular only moderately increases conduction velocity. Higher rates are required to restore conduction velocity to at least half of its normal value. At transduction rates up to 0.4, the risk of conduction block is low. However, at very low levels of coupling in diseased cells, higher levels of viral transduction may be associated with a greater susceptibility to conduction block.

**Table 1.** Summary showing the proportion of transduced cells required to restore conduction velocity to 50% of normal (positive control represented by the solid black symbols in Figures 3, 4 and 5).

| Transduction efficiency needed to restore conduction velocity to 50% of normal |  |  |  |  |
| --- | --- | --- | --- | --- |
| Intercellular coupling level in *PKP2* ACM cells | LR1 model |  | DeW model |  |
|  | wide | narrow | wide | narrow |
| 0.15x normal | 0.6 | 0.6 | 0.6 | 0.7 |
| 0.10x normal | 0.7 | 0.7 | 0.7 | 0.7 |
| 0.05x normal | 0.7 | 0.8* | 0.7 | 0.8 |

\* In the LR1 wide strand model at this level of coupling, conduction blocks occurred at transduction levels of 0.4–0.9. No block occurred in the other cases.

## Discussion

We are now in the era of gene therapy for complex diseases of the heart muscle. A recent report of a phase 1 trial using AAV9 viral transduction in patients with Danon disease showed encouraging results, both in terms of safety and early evidence of efficacy.^21^ In this trial, funded by a company now enrolling *PKP2* ACM patients for AAV gene therapy,^2^ 7 patients were treated with varying doses of AAV9 and followed for up to 54 months. Relatively modest levels of successful gene transduction were seen on serial endomyocardial biopsies.^21^ Most biopsies showed that <25% or 25-50% of ventricular myocytes were transduced. In 3 cases, the proportion of transduced cells declined over time, whereas it apparently increased in 2 cases.^21^ Not surprisingly, a higher proportion of transduced myocytes occurred in patients receiving the highest dose of AAV9.

The results of the recent Danon disease trial are directly relevant to our analysis of potential effects of gene therapy in patients with *PKP2* ACM. For example, our simulations showed only modest increases in conduction velocity when up to 40% of myocytes were transduced. This suggests that a significant increase in ventricular conduction velocity will not occur in *PKP2* ACM patients undergoing gene therapy. However, conduction was mostly safe under conditions simulating effects of *PKP2* ACM gene therapy and conduction block occurred only at extremely reduced levels of intercellular coupling. This was because at most coupling levels, intercellular coupling was still sufficient to allow electrotonic current to pass through the diseased cells, and current was not being absorbed by these cells. Thus, the degree of cellular transduction likely to be achieved in current phase I clinical trials^1-3^ will probably not pose a safety concern. Nevertheless, the anticipated levels of transduction may not increase conduction velocity enough to diminish the risk of slow conduction and reentrant arrhythmias.

Our study was focused solely on the effects of correcting reduced electrical coupling and I_Na_ on ventricular conduction in *PKP2* ACM. Our simulations did not incorporate other features likely to be encountered in ACM patients undergoing gene therapy including the presence of complex patterns of fibrofatty scar tissue (mainly in the right ventricle), and immune signaling related both to the gene therapy itself and the underlying pathophysiology of ACM. Moreover, we did not incorporate into our model the decreased levels of inward rectifier potassium current, I_K1,_ observed in our previous study.^10^ Simulations in that study showed that reduced I_K1_ did not produced major changes in conduction velocity compared to changes in I_Na_ or gap junctional coupling. Although decreased I_K1_ may cause depolarization of resting membranes, heterogeneities in resting membrane potential following gene therapy would be minimized by electrotonic charge redistribution. While we acknowledge these limitations, including factors such as fibrosis and effects of inflammation in computer simulations would have vastly increased the complexity of the models. Furthermore, as these features probably vary widely among PKP2 ACM patients, modeling them would be challenging. Nevertheless, it is plausible to speculate on the potential effects of fibrosis and inflammation in the setting of *PKP2* ACM gene therapy.

For example, the presence of scar tissue might further reduce electrical coupling and create source-sink relationships that could promote reentrant arrhythmogenesis. In this regard, we observed conduction block in the LR1 narrow strand model (Figure 5) under conditions of very low coupling (0.05 of normal) and high levels of transduction (0.4-0.9). No block occurred under these conditions in the wide LR1 strand, because wave fronts have more pathways around sites of block in wider structures. However, the possibility of very low coupling in a narrow bundle of myocytes is not completely unrealistic in hearts of ACM patients with active disease; such narrow bundles may run through areas of fibrosis. Whether such conditions actually occur in *PKP2* ACM patients is unclear, although ECGI studies in ACM patients including those with pathogenic variants in *PKP2* suggest they might.^14^ If so, this raises at least some concern about outcomes following higher levels of transduction.

Inflammation could also be a major determinant of outcomes following gene therapy in PKP2 ACM patients. An immune response to AAV gene therapy itself is well documented and some apparently immune-mediated injury occurred in some patients in the Danon disease trial.^21^ While such responses are usually mild and transient, they could promote adverse effects in patients with a highly arrhythmogenic heart muscle disease such as *PKP2* ACM, especially in view of the known effects of inflammatory mediators on cardiac ion channels and action potentials.^22^ Furthermore, ACM appears to be a chronic inflammatory disease in which innate immune signaling occurs in cardiac myocytes that express ACM variants.^23,24^ Persistent immune activation in ACM cardiac myocytes drives myocardial injury and arrhythmias in part by mobilizing pro-inflammatory myocytes to the heart.^23,24^ While successful gene therapy may shut down immune signaling in transduced cells, non-transduced myocytes will presumably continue to mount an innate immune response and recruit injurious inflammatory cells to the heart.

In conclusion, outcomes following gene therapy in *PKP2* ACM patients may be less favorable than in patients with other forms of heart disease currently being treated with gene therapy such as Danon disease or Friedrich’s ataxia. This stems from basic disease mechanisms in ACM in which arrhythmias result from abnormal tissue electrophysiology (and not solely from abnormal cell electrophysiology) and myocardial injury is related at least in part to cell autonomous inflammatory responses and actions of inflammatory cells mobilized to the heart by diseased myocytes. Thus, while transduced cells may become normalized, non-transduced cells in *PKP2* ACM can continue to exert deleterious effects on tissue electrophysiology and injure neighboring cells that have been corrected by gene therapy.

## Nonstandard Abbreviations and Acronyms

ACM: arrhythmogenic cardiomyopathy
*PKP2*: the human gene for the desmosomal protein plakophilin-2
AAV: adeno-associated virus
Cx43: the gap junction protein connexin-43
Na_v_1.5: the alpha subunit of the cardiac voltage-gated sodium channel
LR1: the Luo_Rudy phase 1 model
DeW: the De Waard et al. model
WT: wildtype
I_Na_: the voltage-gated cardiac sodium current
I_K1_: the inwardly rectifying potassium current

## Acknowledgments/Sources of Funding

This study was supported by the Swiss National Science Foundation (grant 320030-227774 to JPK) and by a grant from the US National Institutes of Health (grant R01-HL148348 to JES).

